## Supplemental Figures and table 1 for "The cardiopharyngeal mesoderm contributes to lymphatic vessel development"

**Supplemental Figure 1**

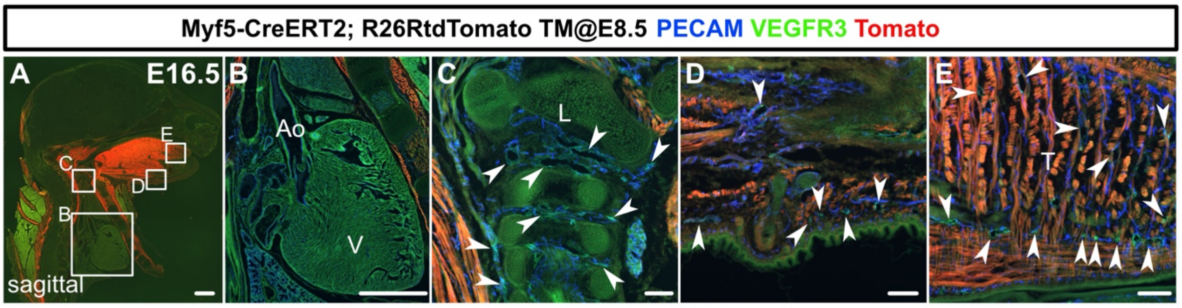


**Supplemental Figure 1. Myf5⁺ lineages do not generate LECs in the cranial or cardiac regions**

(**A-E**) Sagittal sections of Myf5-CreERT2; R26RtdTomato embryos, in which PECAM, tdTomato, and VEGFR3 were labeled at E16.5, are shown. Tamoxifen was administered at E8.5. There were no tdTomato$^{+}$/VEGFR3$^{+}$ lymphatic vessels in or around the heart (**B**), the larynx (**C**), the skin of the lower jaw (**D**), or the tongue (**E**) (white arrowheads indicate tdTomato$^{-}$/VEGFR3$^{+}$ lymphatic vessels). Scale bars, 100 μm **(C-E)**, 1 mm **(A, B)**

**Supplemental Figure 2**

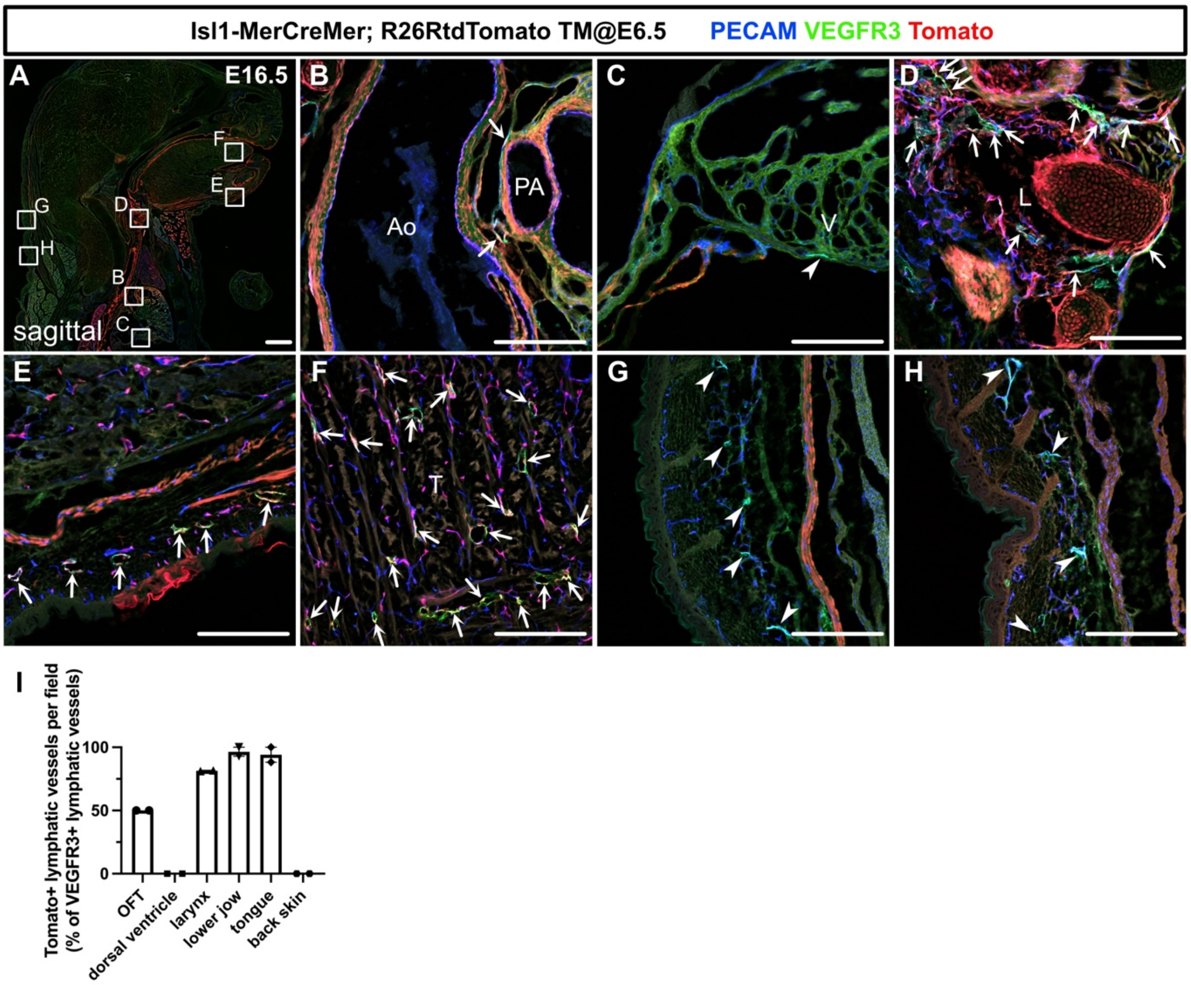


**Supplemental Figure 2. The Isl1$^{+}$ CPM broadly contributes to cranial and cardiac lymphatic vessels**

(**A-H**) Sagittal sections of Isl1-MerCreMer; R26RtdTomato embryos, in which PECAM, tdTomato, and VEGFR3 were labeled at E16.5, are shown. Tamoxifen was administered at E6.5. (**B, D-F**) tdTomato$^{+}$/VEGFR3$^{+}$ lymphatic vessels were observed in and around the cardiac outflow tracts (**B**), the larynx (**D**), the skin of the lower jaw (**E**), and the tongue (**F**) (n=2). (**C, G, and H**) tdTomato$^{+}$/VEGFR3$^{+}$ lymphatic vessels were not observed on the dorsal side of the ventricles (**C**) or in the back skin (**G and H**) (white arrowheads) (n=2). (**I**) The results of a quantitative analysis of the percentage of tdTomato$^{+}$/VEGFR3$^{+}$ lymphatic vessels among all VEGFR3$^{+}$ lymphatic vessels. Each dot represents a value obtained from one sample. Ao, aorta; PA, pulmonary artery; V, ventricle; L, larynx; T, tongue; Scale bars, 100 µm **(B-H)**, 1 mm **(A)**

**Supplemental Figure 3**

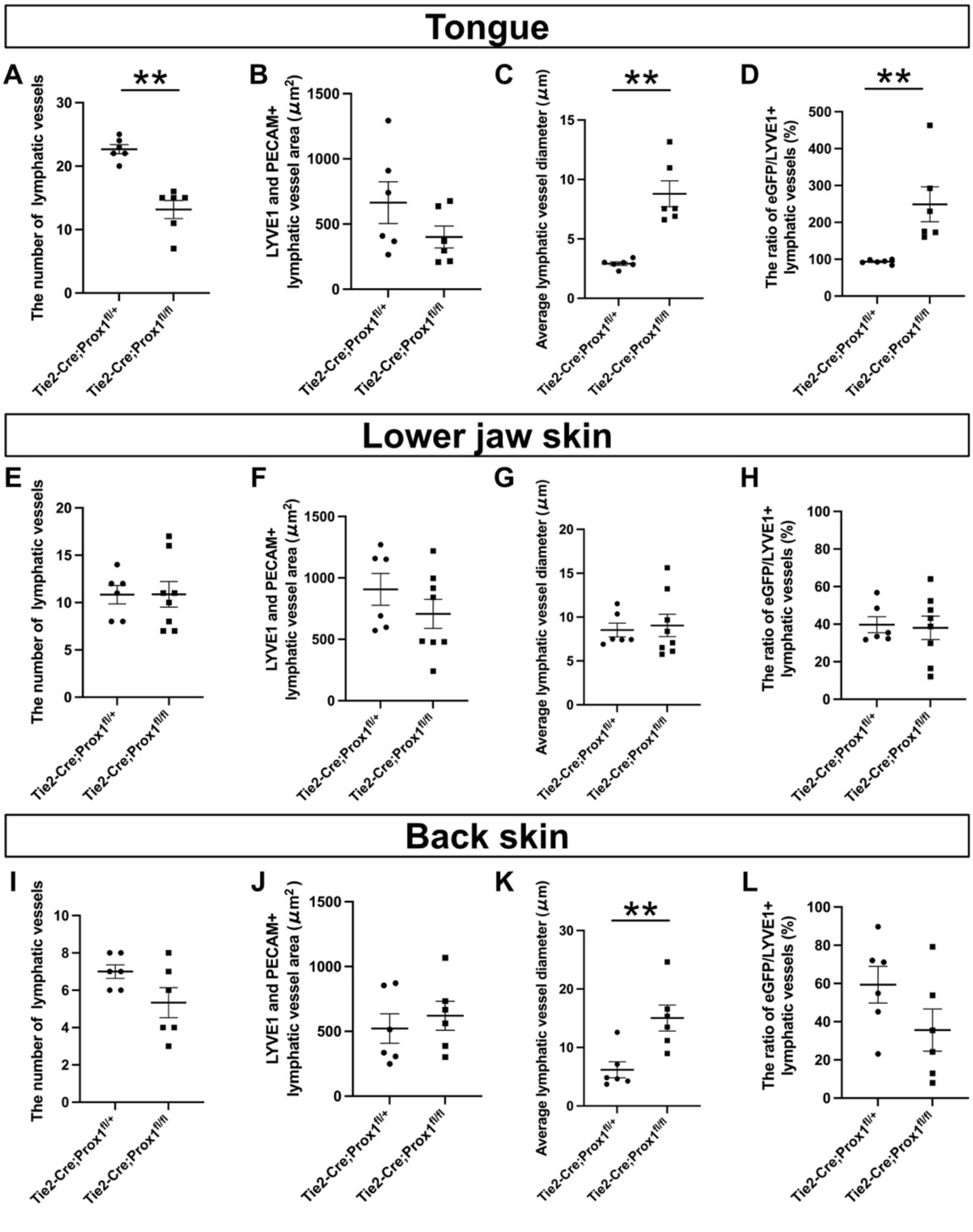


**Supplemental Figure 3. Lymphatic vessel phenotypic differences between Tie2-Cre;Prox1$^{fl/+}$ and Tie2-Cre;Prox1$^{fl/fl}$ embryos**

(**A-L**) The results of quantitative analyses of lymphatic vessel phenotypes in the tongue (**A-D**), the skin of the lower jaw (**E-H**), and back skin (**I-L**) in Tie2-Cre;Prox1$^{fl/+}$ and Tie2-Cre;Prox1$^{fl/fl}$ embryos at E16.5 are shown. All results are expressed as the mean±SEM, and statistical analyses were performed using the non-parametric Mann-Whitney *u*-test. Each dot represents a value obtained from one sample. ***P*<0.01

| | Forward(5'-3') | Reverse(5'-3') |
|---|---|---|
| R26R-eYFP KI | AAAGTCGCTCTGAGTTGTTAT | AAGACCGCGAAGACTTTGTC |
| R26R-eYFP WT | AAAGTCGCTCTGAGTTGTTAT | GGAGCGGGAGAAATGGATATG |
| Rosa26-tdTomato KI | GGCATTAAAGCATATCC | CTGTTCCTGTACGGCATGG |
| Rosa26-tdTomato KI | AAGGGAGCTGCAGTGGAGTA | CCGAAAATCTGTGGGAAGTC |
| Cre | ACATGTTCAGGGATCGCCAG | TAACCAGTGAAACAGCATTGC |
| Prox1flox | CAGCCCTTTTGTTCTGTTGGCCAG | GCAGATGCTGTCCCTACCGTCC |

**Table 1. Primers used for genotyping.**
